# Transcriptome analysis in a humanized mouse model of familial dysautonomia reveals tissue-specific gene expression disruption in the peripheral nervous system

**DOI:** 10.1101/2023.09.28.559870

**Authors:** Ricardo Harripaul, Elisabetta Morini, Monica Salani, Emily Logan, Emily Kirchner, Jessica Bolduc, Anil Chekuri, Benjamin Currall, Rachita Yadav, Serkan Erdin, Michael E. Talkowski, Dadi Gao, Susan Slaugenhaupt

## Abstract

Familial dysautonomia (FD) is a rare recessive neurodevelopmental disease caused by a splice mutation in the Elongator acetyltransferase complex subunit 1 (*ELP1*) gene. This mutation results in a tissue-specific reduction of ELP1 protein, with the lowest levels in the central and peripheral nervous systems (CNS and PNS, respectively). FD patients exhibit complex neurological phenotypes due to the loss of sensory and autonomic neurons. Disease symptoms include decreased pain and temperature perception, impaired or absent myotatic reflexes, proprioceptive ataxia, and progressive retinal degeneration. While the involvement of the PNS in FD pathogenesis has been clearly recognized, the underlying mechanisms responsible for the preferential neuronal loss remain unknown. In this study, we aimed to elucidate the molecular mechanisms underlying FD by conducting a comprehensive transcriptome analysis of neuronal tissues from the phenotypic mouse model *TgFD9*; *Elp1*^Δ*20/flox*^. This mouse recapitulates the same tissue-specific *ELP1* mis-splicing observed in patients while modeling many of the disease manifestations. Comparison of FD and control transcriptomes from dorsal root ganglion (DRG), trigeminal ganglion (TG), medulla (MED), cortex, and spinal cord (SC) showed significantly more differentially expressed genes (DEGs) in the PNS than the CNS. We then identified genes that were tightly co-expressed and functionally dependent on the level of full-length *ELP1* transcript. These genes, defined as *ELP1* dose-responsive genes, were combined with the DEGs to generate tissue-specific dysregulated FD signature genes and networks. Within the PNS networks, we observed direct connections between Elp1 and genes involved in tRNA synthesis and genes related to amine metabolism and synaptic signaling. Importantly, transcriptomic dysregulation in PNS tissues exhibited enrichment for neuronal subtype markers associated with peptidergic nociceptors and myelinated sensory neurons, which are known to be affected in FD. In summary, this study has identified critical tissue-specific gene networks underlying the etiology of FD and provides new insights into the molecular basis of the disease.

## Introduction

Familial dysautonomia (FD) is a rare neurodevelopmental disorder that affects both the peripheral nervous system (PNS) and the central nervous system (CNS) (1–5). The underlying cause of FD is an intronic splice-site mutation in the Elongator acetyltransferase complex subunit 1 gene (*ELP1*, previously known as *IKBKAP*) that results in the tissue-specific skipping of exon 20 (6–10). The nervous system expresses the lowest amount of full-length *ELP1* transcript and protein (11).

*ELP1* encodes subunit 1 of the Elongator Acetyltransferase Complex, which is a highly conserved six-subunit complex (12–17). This complex is involved in numerous cellular functions, including exocytosis, cytoskeletal organization, axonal transport, cellular adhesion, cellular migration of cortical neurons, tRNA modification, and transcriptional elongation (18–31). Through its histone acetyltransferase activity, ELP1 plays a significant role in transcriptional elongation and chromatin organization (13, 32–34),(16, 20). ELP1 also modulates translational efficiency with a biased usage of AA and AG-ending codons, through wobble uridine tRNA modifications (35–38).

FD patients exhibit a range of neurological symptoms that manifest from birth and worsen over time, including diminished pain and temperature sensation, visual loss, kyphoscoliosis, proprioceptive ataxia, and difficulty regulating body temperature and blood pressure (6, 9, 39–43). Loss of sensory neurons in the dorsal root ganglion (DRG), including nociceptors and proprioceptors, is a prominent feature of FD (44–46). Nociceptors are specialized sensory neurons that detect and transmit signals related to pain and temperature perception. In FD patients, there is a diminished ability to perceive pain and temperature, which can lead to insensitivity to potentially harmful stimuli (47). Proprioceptors are sensory neurons responsible for detecting body position and movement and their loss in FD results in proprioceptive ataxia, causing difficulties in coordinating movements and maintaining balance (47, 48). The loss of both nociceptors and proprioceptors contributes to the complex neurological symptoms observed in FD patients.

Mouse models of FD have contributed significantly to our understanding of the role of ELP1 in neural development and function. Multiple studies have provided evidence supporting the crucial role of ELP1 in maintaining neuronal survival and tissue innervation (49–51). The *Elp1* KO mouse provided the first insights into the role of Elp1 in transcriptional elongation and gene expression regulation despite leading to lethality at the mid-gastrulation stage (16). RNA-seq transcriptomic profiling of mouse embryos expressing increasing levels of human ELP1 revealed dysregulation of genes essential to early-stage nervous system development and to the identification of a set of co-expressed genes whose expression highly correlated with the level of ELP1 (29). These ELP1 dose-responsive genes were enriched for axon and cell projection formation which supports the role of ELP1 in the expression of genes important for target tissue innervation and is consistent with the innervation failure observed in FD (29). A phenotypic FD mouse model was generated by introducing the human *ELP1* transgene carrying the FD major splice mutation (*TgFD9*) into a hypomorphic *Elp1^Δ20/Flox^* mouse (*44*). This humanized mouse mimics the tissue-specific mis-splicing seen in FD patients, as well as many phenotypic characteristics of the human disease (52, 53).

While the PNS is known to be significantly affected in FD, the specific gene networks responsible for this disruption have not been identified. In the current study, we uncovered putative, tissue-specific, and convergent molecular mechanisms underlying FD by analyzing the transcriptomes of several neuronal tissues (Figure 1). We collected DRG and trigeminal ganglion (TG) as representative PNS tissues, and cortex, medulla (MED), and spinal cord (SC) as representative CNS tissues, from both control and FD phenotypic mice (Figure 1A) (52). To unravel tissue-specific transcriptomic dysregulation, we identified differentially expressed genes (DEGs) and ELP1 dose-responsive genes (Figure 1A). We then constructed FD-dysregulated gene networks from these transcriptional signatures based on their known protein-protein interactions (Figure 1B). Finally, we compared FD signature genes to determine functional convergence across tissues in FD (Figure 1C). This comprehensive transcriptome analysis provides valuable insights into the regulatory mechanisms underlying FD pathogenesis and sheds light on the shared dysregulation observed in FD PNS tissue.

**Figure 1.**
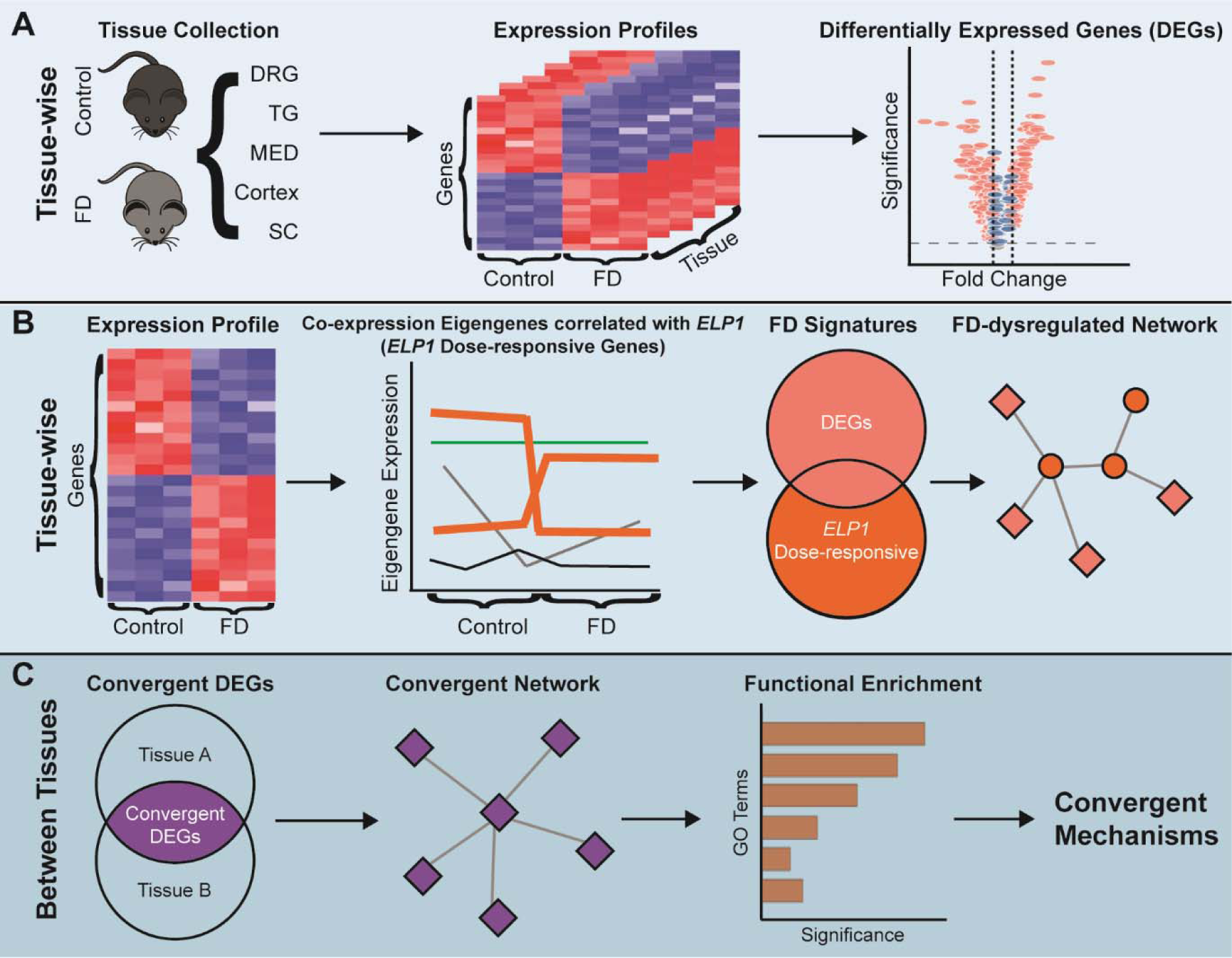
Experiment Design and Analysis Workflow. **(A)** Tissue-specific DEG analyses to reveal the most dominant influence of ELP1 reduction. **(B)** Assembly of FD gene signatures and the disrupted gene network by integrating DEGs and ELP1 dose-responsive genes. **(C)** Identification of convergent disease mechanisms of FD across tissues using shared DEGs.

## Results

### ELP1 reduction leads to tissue-specific transcriptome changes

To uncover the tissue-specific molecular alterations underlying FD, we conducted transcriptome analysis in DRG, TG, cortex, MED, and SC from three-month-old, humanized FD-phenotypic mice (52). We first measured the expression levels of full-length *ELP1* transcript in each tissue and found a significant downregulation of full-length transcript in all five mouse neuronal tissues (Figure 2A). In mouse tissues, the expression of FD full-length *ELP1* transcript compared to controls was 30.97% in DRG, 39.51% in TG, 44.56% in MED, 54.45% in Cortex, and 44.62% in SC. Next, we explored transcriptional ‘signatures’ representing the most significant transcriptional changes across tissues following *ELP1* reduction by performing DEG analyses and gating results on those with false discovery rate (FDR) less than 0.1 and fold changes (FCs) either greater than 1.2 (i.e. upregulated) or less than 0.8 (i.e. downregulated), compared to controls (see Methods). Using this approach, we observed 148 DEGs (FDR < 0.1) in DRG, 194 DEGs in TG, 65 DEGs in MED, 19 DEGs in SC and 59 DEGs in cortex (Figure 2B, Supplementary Figure S1A, Supplementary Table S1), demonstrating significantly higher dysregulation in the PNS tissues. As expected, *ELP1* was the most downregulated gene in all five tissues. The strongest increase in expression was observed with *Fev* (alias *Pet1*) in the DRG, TG, MED, and SC. *Fev* is a transcription factor known to play a crucial role in the differentiation and functional maturation of serotonergic neurons and displayed a 49-fold and 37-fold increase in expression in the DRG and TG respectively (Figure 2B, Supplementary Table S1)(54–59). In DRG and TG, we also observed upregulation of *Th* (1.66-fold in DRG and 2.14-fold in TG), which encodes a tyrosine hydroxylase and serves as a marker for dopaminergic neurons (Figure 2B, Supplementary Table S1).

**Figure 2.**
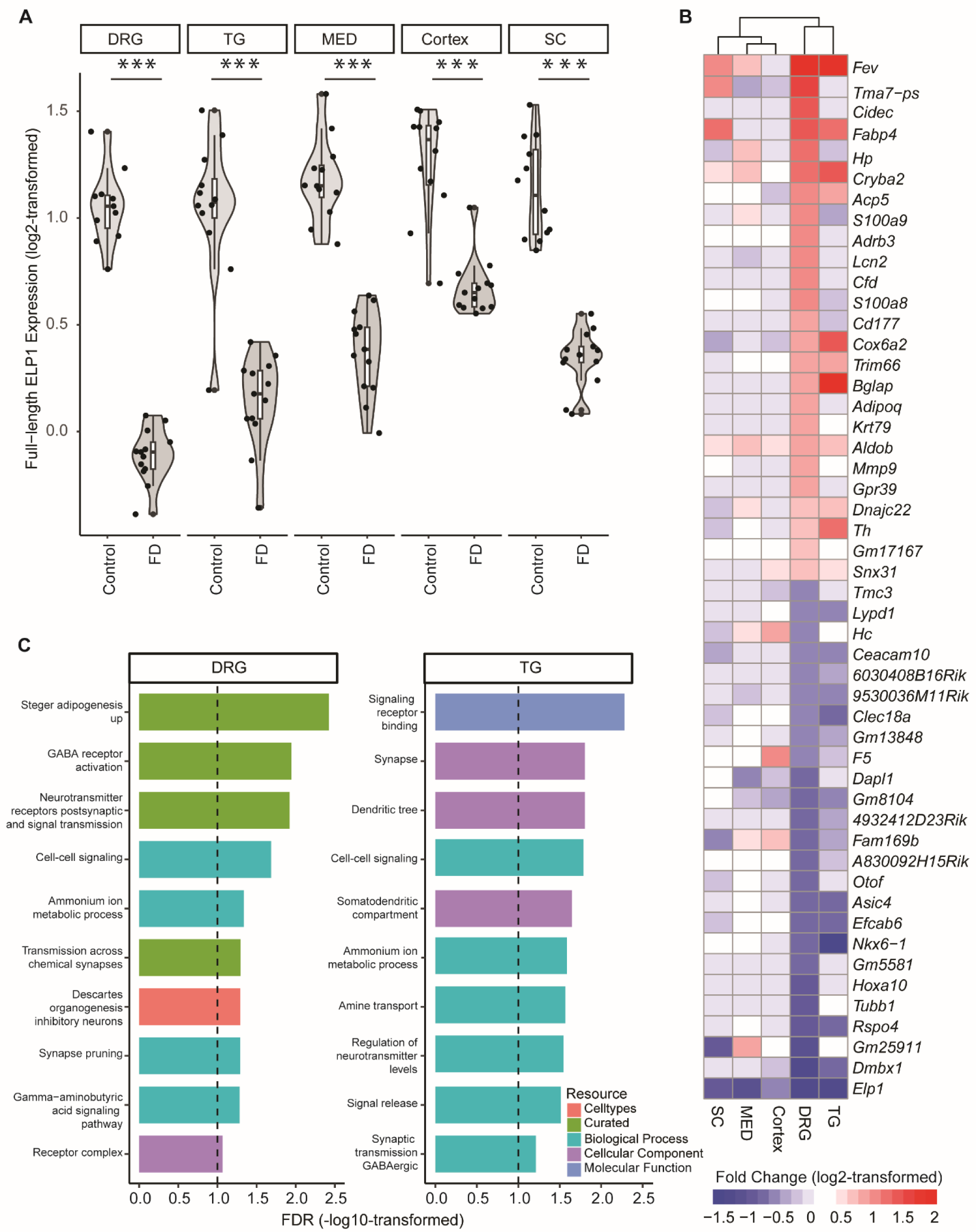
Tissue-specific DEGs and their functional enrichment. **(A)** The violin plot demonstrates the summed expression of full-length human ELP1 and mouse Elp1 in counts per million (CPM, log2-transformed) per tissue. The shape represents the distribution of expression values with individual points representing the actual CPM data points. The line in the middle of the box inside the violin distribution represents the median value with the upper and lower hinges of the box representing the first and third quartiles respectively. A t-test was used to calculate the p-value of the difference between FD and Control, followed by FDR correction. The ‘***’ indicates FDR < 0.001. **(B)** The heatmap represents the 25 most upregulated and downregulated genes between FD and Control across five tissues. Each row represents a gene, and each column represents a tissue. The red color domain represents upregulation between FD and Control while the blue color domain represents downregulation, where the expression changes are measured by log2-transformed fold changes. The deeper the color, the stronger the fold changes. The range was set to [-1.5, 2]. Values beyond this range were clipped to either -1.5 or 2, whichever is closer. The rows are ordered, from top to bottom, by the greatest fold change across the five tissues. **(C)** Bar plot that represents the functional enrichment based on DEGs in DRG and TG, respectively. Note, only 10 significant terms (FDR < 0.1) are selected to display per tissue. The x-axis represents the enrichment significance in -log10-transformed FDR while the y-axis represents the selected significant terms. The bar colors indicate the resources of GO. The vertical black dashed line represents an FDR of 0.1.

We subsequently conducted functional enrichment analyses on the DEGs for each tissue using gene ontology (GO) to identify the pathways that were significantly enriched for DEGs in each FD neuronal tissue (see Methods, Supplementary Table S2). In DRG and TG, which exhibited the most pronounced gene expression dysregulation, we found DEGs were enriched in multiple GO terms associated with synaptic signaling and amine-related metabolic processes at FDR < 0.1 (Figure 2C). Conversely, no significant enrichment was observed in CNS (Supplementary Figure S1B).

The substantial number of observed DEGs, coupled with the significant enrichment of functional terms in the PNS, aligns with the observation of a drastic reduction (average ∼65%) in full-length *ELP1* transcript within the PNS tissues. The dramatic ELP1-dependent gene dysregulation in the PNS is consistent with the significant neuronal loss observed in DRG from FD patients (41). The fact that an average 52% decrease in ELP1 in the CNS does not result in dramatic gene dysregulation underscores different tissue-specific sensitivity to ELP1 reduction.

### Dose-responsive genes create a connection between the DEGs specific to each tissue and ELP1 reduction

Although there were significant expression alterations (DEGs) between FD and control tissues, we did not observe a direct connection to Elp1 in the mouse protein-protein interaction network (PPI). Therefore, we sought to identify ELP1 dose-responsive genes, which are defined as genes that display co-expression and tight correlation with the level of full-length *ELP1* transcript (29). These genes are highly sensitive to ELP1 dosage even though their expression does not significantly change between FD and control. To identify the ELP1 dose-responsive genes, we adopted a two-step approach as previously described in Morini *et al.*, 2022 (29). Initially, we identified co-expressed gene modules with eigengenes that correlate with full-length *ELP1* expression, followed by filtering for individual genes within these modules that display the strongest correlation (see Methods). Among 641 co-expression modules across five tissues, only seven modules (1.09%) in DRG, TG, and MED showed high correlation with the expression of full-length *ELP1* transcript (Supplementary Figure S2A). In DRG, we identified 156 ELP1 dose-responsive genes, while in TG and MED, we identified 137 and 514 such genes, respectively (Supplementary Figure S2B-H, Supplementary Table S3).

Next, we intersected DEGs and ELP1 dose-responsive genes to generate a set of tissue-specific FD signature genes that showed a strong correlation with ELP1 expression. Using the annotated mouse PPI data, we constructed gene networks to uncover the potentially disrupted molecular pathways in FD (see Methods) (Figure 3A, Supplementary Figures S3A). Interestingly in DRG, TG and MED we observed a significantly higher number of interactions among the FD signature genes compared to what would be expected by chance (PPI enrichment *p*-value < 1.0E-12, Figure 3A, Supplementary Figure S3A). In all three tissues, Elp1 was found to be connected to a network that encompassed at least 48% of the FD signature genes (DRG, 48.21% or 121/251; TG, 49.82% or 138/277, and MED 86.27% or 490/568, Supplementary Table S4). Notably, these networks highly relied on the inclusion of the *ELP1* dose-responsive genes (Figure 3A, Supplementary Figures S3A), as their exclusion resulted in an Elp1 network that contained less than 1% of the DEGs. For instance, in the DRG FD-dysregulated network, three *ELP1* dose-responsive genes, namely *Iars* (FDR = 0.041, fold change = 108%), *Asns* (FDR = 0.015, fold change = 110%), and *Aldh18a1* (FDR = 0.011, fold change = 111%), mediated the interactions between Elp1 and all network DEGs, except for *Hdhd3* and *Pxylp1* (Figure 3A). *Iars* encodes isoleucyl-tRNA synthetase (60), *Asns* encodes asparagine synthetase (61), and *Aldh18a1* encodes pyroline-5-carboxylate synthetase (62), which are all associated with cellular amino acid metabolism. They establish connections between Elp1 and a series of solute carrier 7 (Slc7) family members, among which, Slc7a5 and Slc7a3, are responsible for neuronal amino acid transport across the cell membrane and are DEGs between FD and control (63, 64).

**Figure 3.**
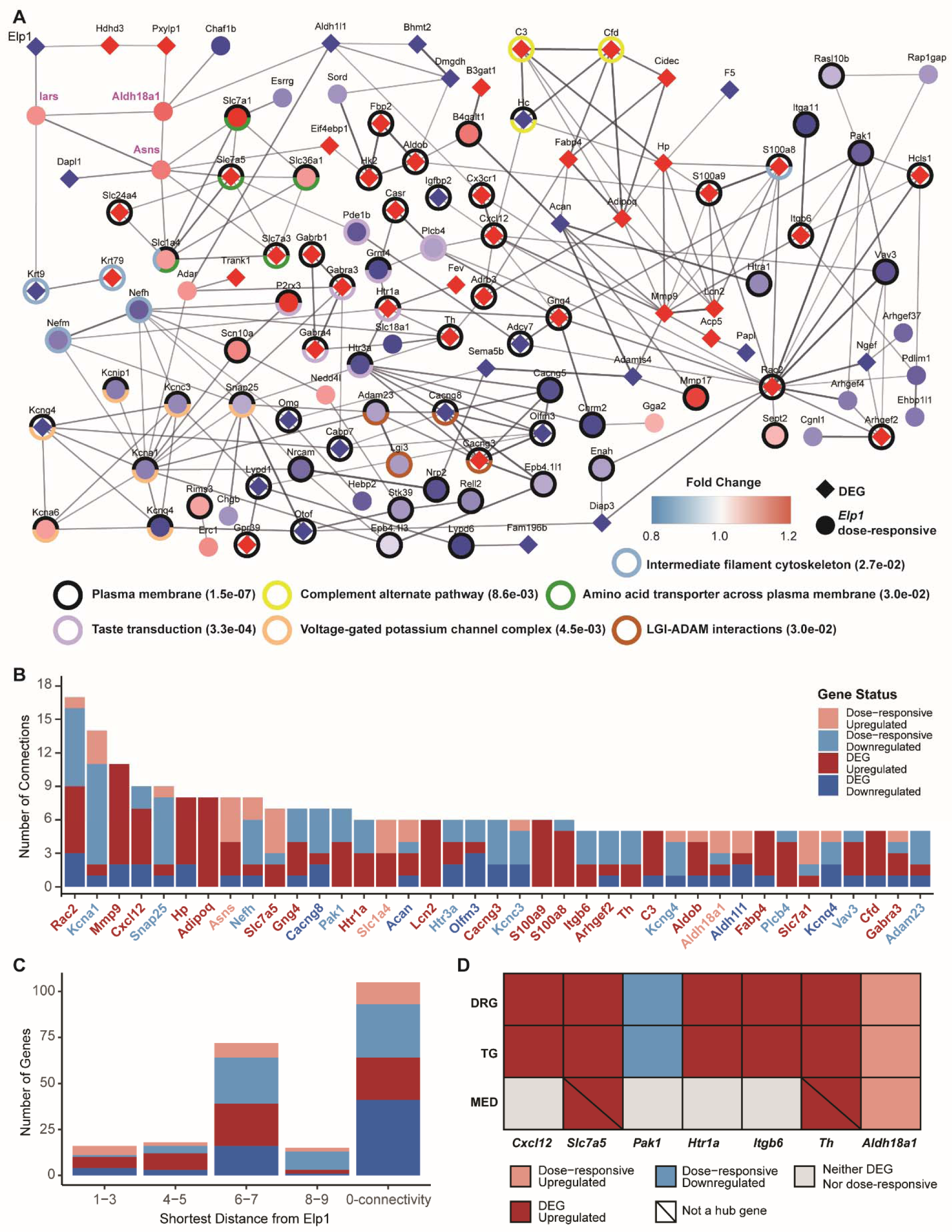
Tissue-specific dysregulated gene network due to ELP1 reduction. **(A)** The dysregulated gene network in DRG. Each node is either a DEG indicated by a diamond shape or an ELP1 dose-responsive gene indicated by a round shape. The colors for the nodes reflect the fold changes in the genes between FD and Control. The red color domain represents upregulation between FD and Control while the blue color domain represents downregulation. The deeper the color, the stronger the fold changes. Each edge represents an interaction between the two connected genes, where only an interaction score of more than 0.4 (default) in String-DB is displayed. The thicker the edge, the higher the interaction score. Only the dysregulated genes with at least one interaction are displayed. The rings outside the nodes represent significant functional enrichment with FDR < 0.1 using all the dysregulated genes (i.e., DEGs and ELP1 dose-responsive genes). The names of three tRNA synthetases next to Elp1 were marked in magenta. The associated functional enrichment terms with the ring colors are given, where the values in the brackets are the enrichment FDRs for the terms. **(B)** The bar plot demonstrates hub genes in DRG ranked by their number of connections to the neighbor genes in the network of panel (A). The x-axis represents the hub gene names, where each gene is colored according to its dysregulation direction and gene category. **(C)** The bar plot demonstrates the number of dysregulated genes in DRG at different distances to ELP1. The x-axis represents the distance of the shortest path to a gene. The genes in the “0-connectivity” distance category refer to those dysregulated genes not displayed in panel (A) because they don’t have any interaction score >= 0.4. The y-axis represents the number of genes at each distance. **(D)** The table demonstrates the shared hub genes across DRG, TG, and MED. The rows represent the tissues while the columns represent the hub genes shared by at least two tissues. The colors of the grids reflect the genes’ categories and dysregulation directions.

The FD-dysregulated network in DRG showed several FD signature genes with five or more connections to their neighbors that serve as “hubs” within the network. It is noteworthy that 53.85% (21/39) of these hub genes were upregulated DEGs, while only 12.82% (5/39) of the hub genes were downregulated DEGs (Figure 3B). To assess their relative contribution to the transcriptomic dysregulation observed in FD DRG, we ranked these hub genes based on their number of connections (Figure 3B). The top five hubs that are DEGs in the network, *Rac2*, *Mmp9*, *Cxcl12*, *Hp,* and *Adipoq*, directly connect to 42.19% (27/64) of the network DEGs in DRG, as well as to each other. Further investigation into the molecular function of these genes and their role in maintaining neuronal health will provide valuable insights into the etiology of FD. We then examined the abundance of DEGs at different distances from Elp1 in the FD-dysregulated network of DRG, which provided insights into the influence of ELP1 reduction on the transcriptome. Interestingly, we found that 28.69% (72/251) of FD signature genes (and 26.35% DEGs) were located six to seven steps away from Elp1 in the DRG dysregulated network (Figure 3C). In contrast, there were fewer FD signature genes (13.55%) within five steps of Elp1. This observation aligns with the placement of the top five hub genes within the network and suggests that the impact of reduced ELP1 levels may be amplified along the FD-dysregulated network in DRG.

The FD signature genes specific to TG showed significant enrichment in synaptic signaling, GABAergic synapse and neurotransmitter pathways (Supplementary Figure S3A, Supplementary Table S5). Similar to DRG, the connection between Elp1 and the rest of the FD signature genes was mediated by an *ELP1* dose-responsive tRNA synthetase, *Cars* (65) (FDR = 0.0011, fold change = 113%, Supplementary Figure S3A). Furthermore, like in DRG, the FD signature genes in TG are located far away from Elp1 (Supplementary Figure S3B), and the majority (71.88%) of TG hub genes are significantly upregulated in FD compared to control samples (Supplementary Figure S3C).

In MED, both the FD signature genes and FD-dysregulated gene network were distinct from those in the PNS tissues. Out of 568 FD signature genes specific to MED, 87.85% (499/568) were *ELP1* dose-responsive genes. The significant number of dose-responsive genes, coupled with the low count of DEGs, suggests a relatively mild impact of ELP1 reduction on the MED transcriptome. The FD signature genes were significantly enriched in the chromatin regulator term (Supplementary Figure S4A, S4C, Supplementary Table S5), which was not observed in the FD signature genes of the PNS tissues. Additionally, these gene signatures were found to be closer to Elp1, with 63.38% (360/568) of MED signature genes located within four steps of Elp1 (Supplementary Figure S4B).

Collectively, our findings indicate that ELP1 dose-responsive genes play a crucial role in mediating the connections between tissue-specific DEGs and Elp1. Furthermore, they nominate highly connected loci in the FD-regulatory gene network, shedding light on important contributors to the molecular etiology of FD.

### Convergence of transcriptomic dysregulation in FD DRG and TG

We compared the hub genes across the three tissue-specific FD networks, and we discovered that seven hubs were shared between DRG and TG, while only one hub was shared by all three tissues (Figure 3D, Supplementary Figure S4B). To identify common and convergent molecular mechanisms underlying FD, we evaluated the extent of similarity between any two transcriptomes in relation to ELP1 reduction. From a differential expression perspective, we observed that DRG and TG shared 44 DEGs (∼ 26%, *p* = 4.55E-54, hypergeometric test) enriched for synaptic signaling, dendrite tree development, and ammonium ion metabolic processes (Figure 4B, Supplementary Table S6). In contrast, there were less than five overlapping DEGs between any two CNS tissues, although the number of overlaps was significantly different from what would be expected by chance (Figure 4A). Among the three CNS tissues, only three DEGs, including *ELP1,* were shared (Supplementary Table S1).

**Figure 4.**
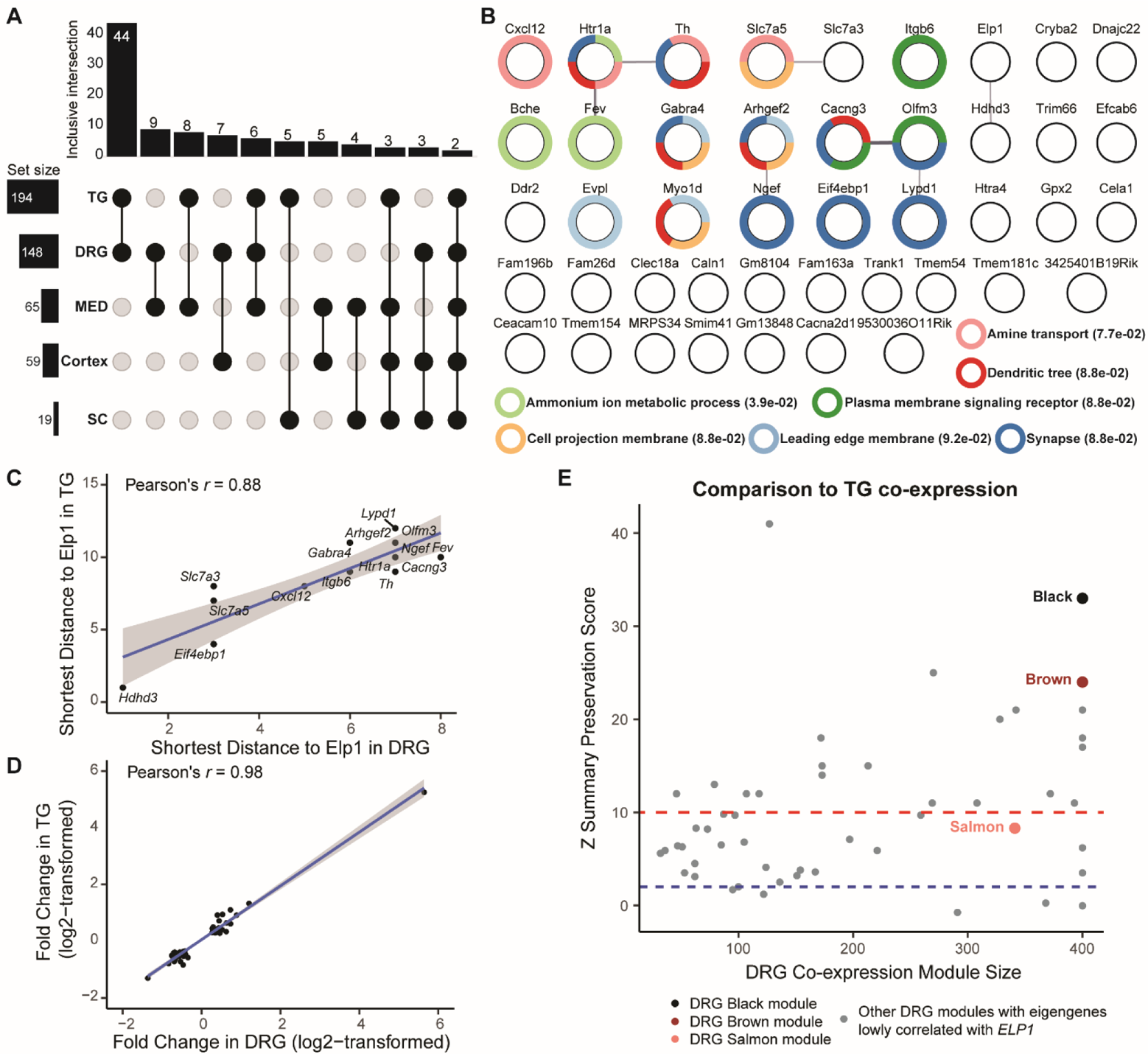
The convergence of transcriptomic dysregulation in the PNS tissues. **(A)** The UPSET plot demonstrates the DEGs overlaps between two out of the five tissues. The x-axis of the bar plot represents overlap comparisons while the y-axis of it represents the number of genes. **(B)** The gene network of PNS convergent DEGs. Each node represents a PNS convergent DEG indicated by a round shape. Each edge represents a potential interaction between the two connected genes, where only an interaction score of more than 0.4 (default) in String-DB is displayed. The thicker the edge, the higher the interaction score. The rings outside the nodes represent significant functional enrichment with FDR < 0.1 of all the PNS convergent DEGs. The associated functional enrichment terms with the ring colors are given, where the values in the brackets are the enrichment FDRs for the terms. **(C)** The scatter plot demonstrates the shortest distances of PNS convergent DEGs from ELP1 in the DRG dysregulated network (x-axis) and the TG dysregulated network (y-axis), respectively. Each dot represents a PNS convergent DEG with non-zero connectivity in both networks. The blue line represents the best-fitted linear regression line while the grey zone around the line represents the 95% confidence intervals. **(D)** The scatter plot demonstrates the log2-transformed fold change of PNS convergent DEGs in DRG (x-axis) and TG (y-axis), respectively. Each dot represents a PNS convergent DEG. The blue line represents the best-fitted linear regression line while the grey zone around the line represents the 95% confidence intervals. **(E)** The scatter plot demonstrates the DRG co-expression modules’ sizes (x-axis) and their similarity to the TG co-expression modules, measured by Z summary preservation scores (y-axis). The score indicates the degree of relatedness of each module to other modules in other co-expression networks. The dots represent the co-expression modules identified in DRG. The three modules whose eigengene highly correlated with the full-length ELP1 expression, namely black, brown, and salmon, are highlighted in their corresponding colors. The blue dashed line indicates a module preservation score of 2 below which the preservation is not considered strong, while the red dashed line represents a module preservation score of 10 above which the preservation is considered very strong.

From a quantitative perspective, the level of transcriptome disruption in FD DRG and TG was found to be similar. The PNS convergent DEGs, when connected to Elp1 in both DRG- and TG-specific FD-dysregulated gene networks, exhibited a significantly similar distribution (Pearson correlation coefficient = 0.88, *p* < 2.2E-16) and were relatively distant from Elp1 (Figure 4C). Additionally, the magnitude of dysregulation of these 44 PNS convergent DEGs was nearly identical, as indicated by a high Pearson correlation coefficient of 0.98 (*p* < 2.2E-16) for the fold change correlation between DRG and TG (Figure 4D). We found that 88.5% (46/52) of the co-expression modules in DRG were preserved in TG (Figure 4E). Taken together, these findings suggest convergent dysfunction in the two PNS tissues in FD.

### PNS convergent DEGs show association with specific neuronal subtypes in FD

To gain a deeper understanding of how PNS convergent DEGs contribute to FD etiology, we hypothesized that the observed dysregulation might be driven by specific neuronal subtypes unique to the PNS. To investigate this, we analyzed publicly available single-cell RNA sequencing (scRNA-seq) data from wildtype mouse DRG (66, 67) and TG (68). By combining neuronal subtype markers provided in these studies with the novel markers identified through our analyses (Supplementary Figure S5A-C, Supplementary Table S7, see Methods), we discovered significant overlaps with the DRG, TG, and PNS convergent DEGs (Figure 5A). These sets of DEGs were enriched for specific neuronal subtype markers. Notably, we observed enrichment of markers for peptidergic nociceptors in DRG and TG, myelinated sensory neurons, and Th+ neurons in DRG, and c-fiber mechanoreceptors and cold nociceptors in TG (all FDRs < 0.07, hypergeometric test). Furthermore, out of 44 PNS convergent DEGs, 15 were identified as neuronal subtype markers (Figure 5B), with 11 of them being markers for peptidergic nociceptors and myelinated sensory neurons (69). These findings suggest that these two neuronal subtypes might be particularly susceptible to ELP1 reduction in the PNS.

**Figure 5.**
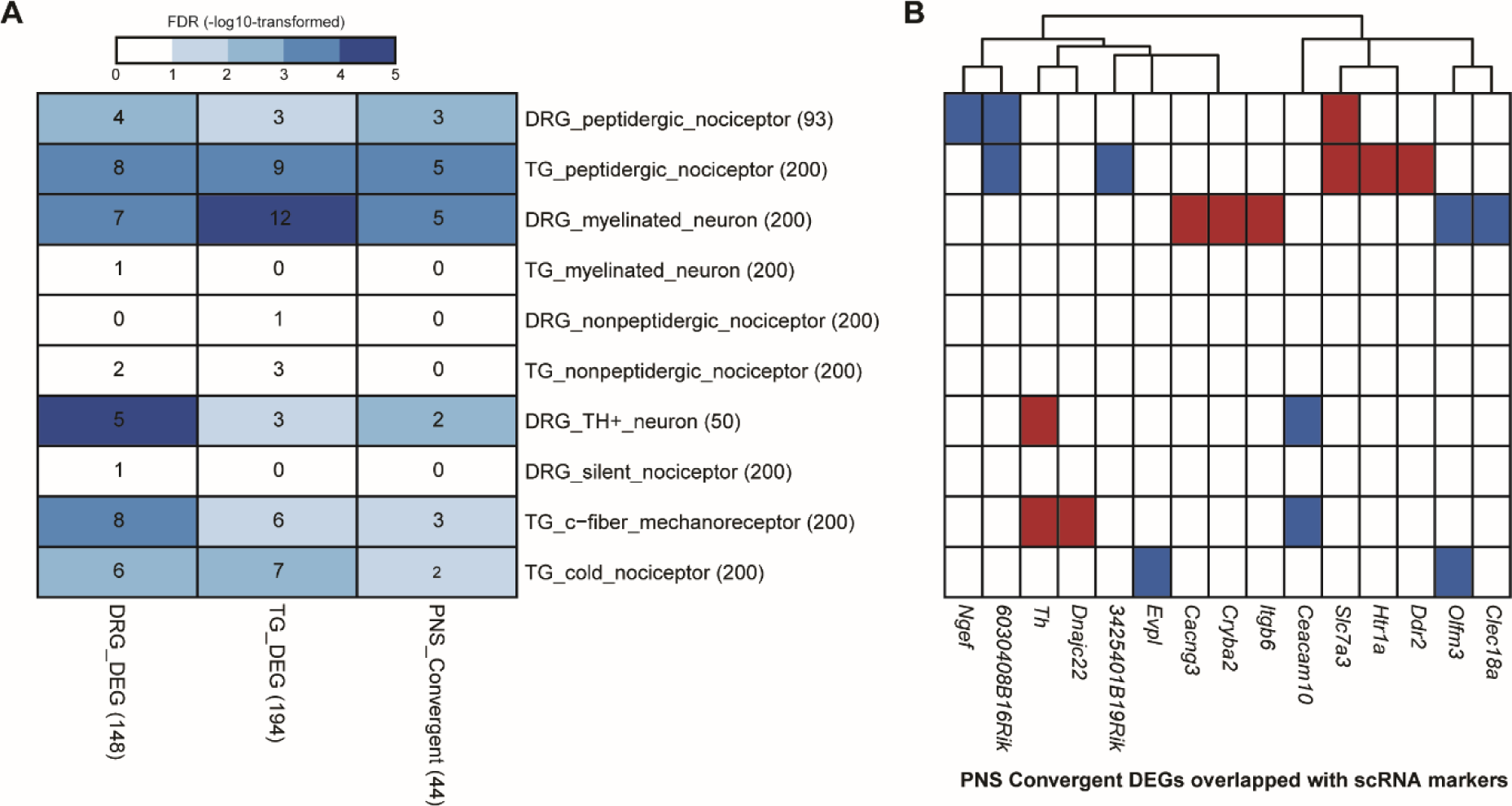
The potential transcriptomic dysregulation at neuronal subtype levels in the PNS tissues. **(A)** The heatmap demonstrates the overlap between DEGs identified from this study (columns) and the neuronal subtype markers identified from publicly available scRNA datasets for mouse DRG and TG. The numbers in the brackets after the row or column names indicate the number of genes in each category. The numbers in the grids indicate the number of gene overlaps between the two categories. The grid colors reflect the significance of overlap according to the hypergeometric test followed by FDR correction, in the -log10-transformed scale. The deeper the color, the more significant the overlap. The grids in white mean no significant overlap where FDR >=0.1. **(B)** The overlaps between the PNS convergent DEGs (columns) and the neuronal subtype markers identified from publicly available scRNA datasets for mouse DRG and TG (rows, the same as panel (A)). The colors of the grids reflect the genes’ dysregulation directions between FD and Control, where red is for upregulation while blue is for downregulation.

Collectively, our results provide novel insights into the dysregulation of peripheral nervous system gene expression in FD. Tissue-specific analyses revealed a greater impact of ELP1 reduction on PNS tissues compared to CNS tissues, as evidence by the number of DEGs. The FD-dysregulated gene networks showed upregulated hub genes that were significantly enriched in the PNS tissues. Cross-tissue comparisons further highlighted convergent mechanisms of disrupted synaptic signaling and amine-related metabolic processes in the PNS tissues, while such convergence was not observed across CNS tissues.

## Discussion

FD is a progressive neurodegenerative disease that manifests in various debilitating symptoms including diminished pain and temperature perception, decreased or absent myotatic reflexes, proprioceptive ataxia, and retinal degeneration. Recent studies have provided compelling evidence linking the reduction of Elp1 to sensory neuronal loss and diminished tissue innervation (70–72). However, the intricate molecular mechanisms connecting ELP1 reduction with the phenotypic manifestations of the disease remain largely unknown.

Using a humanized FD phenotypic mouse that recapitulates the same tissue-specific reduction of ELP1 observed in patients, we have conducted a comprehensive transcriptomic study to investigate the disrupted gene expression and pathways underlying FD etiology in disease-relevant neuronal tissues (44). We collected DRG and TG as representative PNS tissues, and cortex, MED, and SC as representative CNS tissues, from both control and FD-phenotypic mice. We found significant differences in the levels of full-length *ELP1* transcripts between PNS and CNS. The PNS tissues, DRG and TG, exhibited lower levels of full-length *ELP1* transcripts (∼ 35% of the control level) when compared to the three CNS tissues, MED, cortex, and SC (∼48% of the control level).

To gain deeper insights into the molecular networks and pathways involved in FD, we assembled a set of FD signature genes by combining tissue-specific DEGs that represented the most prominent transcriptional alterations, and ELP1 dose-responsive genes that exhibited moderate expression changes but they closely correlated with ELP1 levels. The FD signature genes formed interconnected gene networks providing a comprehensive view of how ELP1 reduction impacts the expression of many genes. This study shows that merely using DEGs is not sufficient to build a meaningful PPI network. Adding dose-response genes interconnects the robust signals from DEGs and creates a more interconnected and expansive network. FD is a recessive disease but, unlike most recessive diseases, it does not result from complete loss of a protein but is caused by tissue-specific reduction of ELP1(73). It is reasonable to think that the reduction of ELP1, instead of its complete depletion, might cause moderate transcriptomic changes (e.g. *ELP1* dose-responsive genes) in its immediate downstream genes. These moderate responders could then synergistically lead to more dramatic expression alterations (DEGs) deeper in the molecular network. We indeed observed such patterns in both DRG and TG FD-dysregulated networks.

In the DRG, for instance, the connection of Elp1 with the other DEGs is dependent on the inclusion of the three dose-responsive genes *Iars*, *Aldh18a1* and *Asns* (Figure 3A). These three genes encode synthase proteins. It is well known that the cellular concentrations of tRNA synthase must be precisely regulated and overproduction of them leads to various disorders including neurodegeneration (38, 74). In addition, we observed significant upregulation of amino acid transporter genes *Slc7a5* (alias *Lat1*) and *Slc7a3* (alias *Cat3*). Both tRNA synthesis and amino acid transport are the immediate upstream biological steps of tRNA wobble modification, one of the well-studied functions of *ELP1*(*23, 25, 27, 38, 74*). We acknowledge that these findings are based on the expression profiles identified from an FD-phenotypic mouse model and rely on the annotated mouse PPI network. Further evaluation is needed to determine the extent to which these findings can be translated to FD patients. Transcriptome-wide, we observed striking similarities in the response to ELP1 reduction between DRG and TG. The PNS convergent DEGs in this study were enriched for synaptic signaling and amine transport. This PNS enrichment aligns with the functional enrichment patterns observed in DEGs specific to each tissue. Further exploration of these convergent and tissue-specific DEGs may provide valuable insights into the underlying mechanisms of FD.

Finally, to determine if the observed dysregulation was specific to unique neuronal subtypes in the PNS, we combined the markers identified through our analyses with mouse DRG and TG neuronal subtype markers from publicly available single-cell RNA sequencing (scRNA-seq) data (66, 67),(68). Remarkably, we found significant overlaps between the PNS convergent DEGs and the single-cell markers associated with peptidergic nociceptors and myelinated sensory neurons supporting the hypothesis that certain neuronal subtypes are more susceptible to ELP1 reduction.

Overall, our study provided a comprehensive analysis of the disrupted transcriptomic dynamics in FD from both tissue-specific and cross-tissue perspectives. By examining gene expression patterns, we identified several gene sets that may contribute to the disease manifestations. The FD signature genes in the PNS tissues were found to be enriched in amine-related metabolic processes, which in turn influenced synaptic signaling. Our findings suggest the peptidergic nociceptors and myelinated sensory neurons in the PNS are particularly affected in FD, as evidenced by significant alterations in the expression of their marker genes upon ELP1 reduction. Our study not only provides valuable insights into the intricate molecular mechanisms underlying FD but also have broader implications for other neurological disorders associated with Elongator dysfunction.

## Methods

### Animals

The generation of the *TgFD9* mouse line carrying the human *ELP1* transgene with the NM_003640.5:c.2204+6T>C mutation can be found in Hims et al. (75). Descriptions of the original targeting vector used to generate the *Elp1^flox^* allele and the strategy to generate the *Elp1*^Δ*20*^ allele have been previously published (76, 77).

To generate the experimental *TgFD9; Elp1*^Δ*20/flox*^ mouse, we crossed the *TgFD9* transgenic mouse heterozygous for the *Elp1^flox^* allele (*TgFD9^+/-^*; *Elp1^flox/+^*) with each other. Pups were genotyped to identify those homozygotes for both the *TgFD9* transgene and the *Elp1^flox^* allele (F1: *TgFD9^+/+^; Elp1^flox/flox^*). These animals were then crossed with the mouse line heterozygous for the *Elp1*^Δ*20*^ allele (*E1p1*^Δ*20/+*^) to generate the FD mouse *TgFD9; Elp1*^Δ*20/flox*^ (F2). Controls are littermates of the FD mice that carry the transgene but are phenotypically normal because they express the endogenous *Elp1 gene (TgFD9^+/-^; Elp1^+/+^, TgFD9^+/-^; Elp1 ^flox/+^ or TgFD9^+/-^; Elp1* ^Δ^*^20/+^)*. Both sexes were included in this study. The mice were housed in the animal facility at Massachusetts General Hospital (Boston, MA), provided with access to food and water *ad libitum*, and maintained on a 12-hour light/dark cycle. All experimental protocols were approved by the Institutional Animal Care and Use Committee of the Massachusetts General Hospital and were in accordance with NIH guidelines.

For routine genotyping of progeny, genomic DNA was prepared from tail biopsies, and PCR was carried out using the following primers - forward, 5‘-TGATTGACACAGACTCTGGCCA-3’; reverse, 5‘-CTTTCACTCTGAAATTACAGGAAG-3’-to discriminate the *Elp1* alleles and the primers - forward 5‘-GCCATTGTACTGTTTGCGACT-3’; reverse, 5‘-TGAGTGTCACGATTCTTTCTGC-3’-to detect the *TgFD9* transgene.

### Tissue-Specific RNA-seq

RNA was extracted from DRG, trigeminal ganglion, cortex, medulla, and spinal cord collected from 12 control (6 males and 6 females) and 13 FD (4 males and 9 females) 3-month-old mice. Using the QIAzol Reagent, following the manufacturer’s instructions. RNA-seq libraries were prepared using the Tru-Seq Stranded® mRNA library Prep Kit (Illumina, 20020594) using 100 ng of total RNA as input. Final library concentration was quantified using size distribution by the Agilent 2200 Tape Station and/or qPCR using the Library Quantification Kit (KK4854, Kapa Biosystems). Equimolar amounts of each library were pooled prior to multiplex sequencing. Libraries were 50 basepair paired end sequenced on the Illumina HiSeq 2500 across multiple lanes. The HiSeq Sequencing Control Software was used for real-time image analysis and base calling before CASAVA (version 1.8) was used for FastQ generation.

### RNA-seq Pre-Processing

A custom transcriptome reference was generated by adding the human ELP1 gene (ENSG00000070061) Ensemble Human transcriptome reference GRCh37.75 to the Ensembl Mouse Transcriptome GRCm38.83 as an independent chromosome. RNA-seq reads were mapped to this synthesized transcriptome reference by STAR v2.5.22b allowing only uniquely mapped reads with 5% mismatch (78). Illumina TruSeq reads were trimmed using Trimmomatic (v0.36) with minimal length set to 105 and other default parameters (79).

### Differential Gene Expression Analysis

Gene counts were performed via HTSeq-counts v0.11.2 (80) with ‘-s reverse’ option to be compatible with the Illumina TruSeq library. Genes were further filtered so that only genes whose median expression was no less than 0.1 counts-per-million in at least one genotype were kept for analysis. Raw gene counts were then normalized using sample-wise size factors estimated by the Bioconductor package DESeq2 (v1.34.0)(81). To get the most robust DEGs between the two genotypes of interest (FD and control), surrogate variables unrelated to genotype were first estimated from the normalized counts via the Bioconductor package SVA (v3.42.0, (82)) and then built into a generalized linear model (GLM) together with genotype via DESeq2. Data from both males and females were combined for this analysis. We performed a correlation analysis on several parameters including sex with principal components and surrogate variables. Sex was corrected by surrogate variable analysis and did not correlate with any principal components. Since the phenotypes in FD patients do not exhibit sex differences (83), in this study we aimed at identifying the common disease mechanism regardless of sex.

### Gene Networks According to PPI

These networks were built to include the query gene sets using the “stringApp” (version 2.0.1, (84) in Cytoscape (version 3.9.1) (85). For a tissue-specific FD-dysregulated gene network, the query set consisted of tissue-specific DEGs and ELP1 dose-responsive genes. For the PNS convergent network, the query set consisted of shared DEGs between DRG and TG. To construct each network, the “STRING: protein query” mode was used, with species set as “*Mus musculus*”, confidence (score) cutoff set as 0.4 and maximum additional interactor set as 0.

### Shortest Distance from Elp1

The “Edge Table” from the stringApp network construction was exported. A customized R script was used to convert the pair-wised interactions in the “Edge Table” to an R list object, where the list names represented the nodes (i.e., genes) in the network while the list elements were vectors representing genes connected to each node. Then the shortest distance was calculated via the “shortest_paths” function from an opensource R package “igraph” (version 1.3.5, https://github.com/igraph/igraph).

### GO Analysis

In each analysis, the query gene set was searched against a background gene list consisting of either expressed genes from the same tissue or non-redundant union of expressed genes from multiple tissues where the query set was built from. For functional enrichment of tissue-specific DEGs, the resources of GO were from the Gene Set Enrichment Analysis website (https://www.gsea-msigdb.org/, v. MS1, (86)). For functional enrichment of FD signature genes or PNS convergent DEGs that were used to build interaction networks, the “Functional enrichment” function from the stringApp in Cytoscape was used, with the appropriate background expressed genes as reference. For any two significant functional terms, an overlap score was calculated to reflect their semantic similarity. An overlap score cutoff of either 0.1 or 0.2 was applied on the raw result to remove redundancy in the results.

For all the functional enrichment analyses in this study, complete lists of the results were provided in the supplementary tables.

### Co-expression Module Analysis

Once generalized linear models for differential gene expression were established, the effects from surrogate variables were regressed out from the normalized gene counts to create a cleaned matrix whose variance was mainly due to the genotype difference. Then the R package WGCNA (v1.71) was implemented upon this cleaned matrix of each tissue, identifying genes co-expressed together and grouping them into modules (87). To achieve the best performance, the soft-thresholding power was heuristically selected for each tissue (cortex power = 5, dorsal root ganglion power = 6, medulla power = 5, spinal cord power = 6, trigeminal ganglion power = 7) at the beginning of the WGCNA approach. A signed network was used, and minimal module size was set to 30 and the raw modules were merged with a dis-similarity cut-off of 0.25.

### Correlation between Co-expression Modules’ Eigengenes and the Full-length *ELP1*

#### Expression

The full-length *ELP1* transcript expression was measured as the expression sum of exon 20 (human) and exon 26 (mouse), in the unit of counts per million (CPM). The values across all samples from the same tissue were correlated with each eigengene representing the identified co-expression modules using Pearson correlation. *ELP1* dose-responsive genes were defined as the genes meeting both of the following criteria: 1) the Pearson correlation between their co-expression module eigengene and the full-length *ELP1* had a coefficient no less than 0.8; and 2) the absolute value of the Pearson correlation coefficient between their normalized expression and the module eigengene had a coefficient no less than 0.8.

#### Analysis of Publicly Available scRNA-seq Data

For DRG scRNA-seq, the processed data was downloaded from GEO (GSE59739). Cells with the top and bottom 2.5% of the number of RNA features were filtered out. The normalized counts were what the authors provided(66–68). Dimensionality reduction was first done via multiple correspondence analysis (MCA) using the CelliD package version 1.6.0 (88), followed by uniform manifold approximation and projection (UMAP) using the Seurat package version 4.2.1 (89). Unsupervised clustering was done in the UMAP space using 20-nearest neighbor graph construction with a resolution of 0.2. For each unsupervised cluster, its gene signatures were defined as the top 200 nearest genes to the cluster (i.e., cells) center in the MCA space. The cell-type markers provided by the authors were then compared with the unsupervised cluster signatures. If the overlap was significant under the hypergeometric test, the unsupervised cluster would be marked as the cell-type provided by the authors and the cell-type markers would be replaced by the unsupervised cluster signatures. If the cell-type provided by the authors was found to have no significant overlaps with the markers from the unsupervised clusters, the authors’ cell-types and markers were retained in the final marker list.

For TG scRNA-seq, the raw counts data were downloaded from GEO (GSE197289). Cells with top and bottom 2.5% of the number of RNA features were filtered out. Normalization was done using SCTransform package version 0.3.5 (90), with variance stabilization flavor set to “v2”. Like the DRG processing, the dimension reduction was done by MCA followed by UMAP. The UMAP visual separation already agreed with the cell types provided by the authors. The signature of each cell type was called by the top 200 nearest genes to the cell-type center in the MCA space.

### Statistical Analysis

Wald test was used to estimate the significance of DEGs from the DESeq2 models. Within each tissue, genes with false discovery rate (FDR) < 0.1 and a fold change cut-off was applied (more than 120% for upregulated or less than 80% for downregulated genes) and these genes were then considered as significant. Fisher’s exact test was used for GO analysis where a significant enrichment was defined as FDR < 0.1. A significant correlation throughout this study was defined as Pearson correlation coefficient ≥ 0.8. For overlap significance, hypergeometric test was used and the *p*-value < 0.05 (or FDR < 0.1 when multiple test correction was applicable) was considered as significant. The four values used in the hypergeometric test were the size of gene list A, gene list B, their overlaps, and their non-redundant background genes (e.g., all expressed genes in the transcriptome where list A and B derived from).

## Supporting information

Figure Legends for Main Figures

Supplementary and Supporting Figures with Legends

Differentially Expressed Genes statistics for all tissues

Differentially Expressed Genes - Gene Ontology Enrichment

ELP1 dose-response genes in Ensembl ID form

StringDB Network with Edges using Differentially Expressed Genes and ELP1 dose-response genes

StringDB Network Functional Enrichment using Differentially Expressed Genes and ELP1 dose response genes

Peripheral Nervous System Convergent Differentially Expressed Genes Functional Enrichment

single-cell RNA Markers MCA in DRG and TG

## Data Availability

RNA-seq raw and processed data used in this study is available at GSE230867.

## Acknowledgements

We thank Dr. Alejandra Gonzalez-Duarte and Dr. Horacio Kaufmann of the Dysautonomia Treatment and Evaluation Center at New York University Medical School for their long-standing collaboration and helpful discussions. We are also grateful to Drs. Ioannis Dragatsis and Paula Dietrich at the University of Tennessee for the gift of the Elp1^flox/+^ and Elp1^Δ20/+^ mice and their collaborative effort to generate the FD phenotypic mouse model. This work was supported by National Institutes of Health (NIH) grants (R37NS095640 to S.A.S.).

## Author contributions

**Ricardo S. Harripaul**: Conceptualization, Methodology, Software, Formal analysis, Writing - Original draft. **Elisabetta Morini**: Conceptualization, Methodology, Validation, Formal analysis, Investigation, Writing - Original draft. **Monica Salani**: Investigation, Writing - Review & Editing. **Emily Logan**: Investigation. **Emily G. Kirchner**: Investigation. **Jessica Bolduc**: Investigation. **Anil Chekuri**: Data curation, Investigation. **Benjamin Currall**: Data Curation, Methodology. **Rachita Yadav**: Methodology, Writing – Review & Editing. **Serkan Erdin**: Methodology, Writing - Review & Editing. **Michael E. Talkowski**: Conceptualization, Supervision, Writing - Review & Editing**. Dadi Gao**: Conceptualization, Supervision, Writing - Review & Editing. **Susan Slaugenhaupt**: Conceptualization, Supervision, Writing - Review & Editing, Funding acquisition.

## References

1. Mendoza-Santiesteban CE, Hedges III TR, Norcliffe-Kaufmann L, Warren F, Reddy S, Axelrod FB, et al. Clinical neuro-ophthalmic findings in familial dysautonomia. Journal of neuro-ophthalmology. 2012;32(1):23–6.

2. Mahloudji M, Brunt P, McKusick V. Clinical neurological aspects of familial dysautonomia. Journal of the neurological sciences. 1970;11(4):383–95.

3. Pearson J. Familial dysautonomia (a brief review). Journal of the Autonomic Nervous System. 1979;1(2):119–26.

4. Axelrod FB, Hilz MJ, Berlin D, Yau PL, Javier D, Sweat V, et al. Neuroimaging supports central pathology in familial dysautonomia. Journal of neurology. 2010;257:198–206.

5. Mendoza-Santiesteban CE, Palma J-A, Hedges III TR, Laver NV, Farhat N, Norcliffe-Kaufmann L, et al. Pathological confirmation of optic neuropathy in familial dysautonomia. Journal of Neuropatholgy & Experimental Neurology. 2017;76(3):238–44.

6. Axelrod FB, Nachtigal R, Dancis J. Familial dysautonomia: diagnosis, pathogenesis and management. Adv Pediatr. 1974;21:75–96.

7. Axelrod FB. Familial dysautonomia: a review of the current pharmacological treatments. Expert Opin Pharmacother. 2005;6(4):561–7.

8. Pearson J. Familial dysautonomia (a brief review). J Auton Nerv Syst. 1979;1(2):119–26.

9. Pearson J, Pytel B. Quantitative studies of ciliary and sphenopalatine ganglia in familial dysautonomia. J Neurol Sci. 1978;39(1):123–30.

10. Slaugenhaupt SA, Gusella JF. Familial dysautonomia. Curr Opin Genet Dev. 2002;12(3):307–11.

11. Cuajungco MP, Leyne M, Mull J, Gill SP, Lu W, Zagzag D, et al. Tissue-specific reduction in splicing efficiency of IKBKAP due to the major mutation associated with familial dysautonomia. Am J Hum Genet. 2003;72(3):749–58.

12. Krogan NJ, Greenblatt JF. Characterization of a six-subunit holo-elongator complex required for the regulated expression of a group of genes in Saccharomyces cerevisiae. Mol Cell Biol. 2001;21(23):8203–12.

13. Hawkes NA, Otero G, Winkler GS, Marshall N, Dahmus ME, Krappmann D, et al. Purification and characterization of the human elongator complex. J Biol Chem. 2002;277(4):3047–52.

14. Li F, Lu J, Han Q, Zhang G, Huang B. The Elp3 subunit of human Elongator complex is functionally similar to its counterpart in yeast. Mol Genet Genomics. 2005;273(3):264–72.

15. Chen Z, Zhang H, Jablonowski D, Zhou X, Ren X, Hong X, et al. Mutations in ABO1/ELO2, a subunit of holo-Elongator, increase abscisic acid sensitivity and drought tolerance in Arabidopsis thaliana. Mol Cell Biol. 2006;26(18):6902–12.

16. Chen YT, Hims MM, Shetty RS, Mull J, Liu L, Leyne M, et al. Loss of mouse Ikbkap, a subunit of elongator, leads to transcriptional deficits and embryonic lethality that can be rescued by human IKBKAP. Mol Cell Biol. 2009;29(3):736–44.

17. Kojic M, Wainwright B. The Many Faces of Elongator in Neurodevelopment and Disease. Front Mol Neurosci. 2016;9:115.

18. Rahl PB, Chen CZ, Collins RN. Elp1p, the yeast homolog of the FD disease syndrome protein, negatively regulates exocytosis independently of transcriptional elongation. Mol Cell. 2005;17(6):841–53.

19. Johansen LD, Naumanen T, Knudsen A, Westerlund N, Gromova I, Junttila M, et al. IKAP localizes to membrane ruffles with filamin A and regulates actin cytoskeleton organization and cell migration. J Cell Sci. 2008;121(Pt 6):854–64.

20. Close P, Hawkes N, Cornez I, Creppe C, Lambert CA, Rogister B, et al. Transcription impairment and cell migration defects in elongator-depleted cells: implication for familial dysautonomia. Mol Cell. 2006;22(4):521–31.

21. Tourtellotte WG. Axon Transport and Neuropathy: Relevant Perspectives on the Etiopathogenesis of Familial Dysautonomia. Am J Pathol. 2016;186(3):489–99.

22. Naftelberg S, Abramovitch Z, Gluska S, Yannai S, Joshi Y, Donyo M, et al. Phosphatidylserine Ameliorates Neurodegenerative Symptoms and Enhances Axonal Transport in a Mouse Model of Familial Dysautonomia. PLoS Genet. 2016;12(12):e1006486.

23. Huang B, Johansson MJ, Byström AS. An early step in wobble uridine tRNA modification requires the Elongator complex. Rna. 2005;11(4):424–36.

24. Esberg A, Huang B, Johansson MJ, Byström AS. Elevated levels of two tRNA species bypass the requirement for elongator complex in transcription and exocytosis. Mol Cell. 2006;24(1):139–48.

25. Karlsborn T, Tükenmez H, Chen C, Byström AS. Familial dysautonomia (FD) patients have reduced levels of the modified wobble nucleoside mcm(5)s(2)U in tRNA. Biochem Biophys Res Commun. 2014;454(3):441–5.

26. Yoshida M, Kataoka N, Miyauchi K, Ohe K, Iida K, Yoshida S, et al. Rectifier of aberrant mRNA splicing recovers tRNA modification in familial dysautonomia. Proc Natl Acad Sci U S A. 2015;112(9):2764–9.

27. Bauer F, Hermand D. A coordinated codon-dependent regulation of translation by Elongator. Cell Cycle. 2012;11(24):4524–9.

28. Chen WT, Tseng HY, Jiang CL, Lee CY, Chi P, Chen LY, et al. Elp1 facilitates RAD51-mediated homologous recombination repair via translational regulation. J Biomed Sci. 2021;28(1):81.

29. Morini E, Gao D, Logan EM, Salani M, Krauson AJ, Chekuri A, et al. Developmental regulation of neuronal gene expression by Elongator complex protein 1 dosage. J Genet Genomics. 2022;49(7):654–65.

30. Creppe C, Malinouskaya L, Volvert M-L, Gillard M, Close P, Malaise O, et al. Elongator controls the migration and differentiation of cortical neurons through acetylation of α-tubulin. Cell. 2009;136(3):551–64.

31. Xu Y, Zhou W, Ji Y, Shen J, Zhu X, Yu H, et al. Elongator promotes the migration and invasion of hepatocellular carcinoma cell by the phosphorylation of AKT. International Journal of Biological Sciences. 2018;14(5):518.

32. Otero G, Fellows J, Li Y, de Bizemont T, Dirac AM, Gustafsson CM, et al. Elongator, a multisubunit component of a novel RNA polymerase II holoenzyme for transcriptional elongation. Mol Cell. 1999;3(1):109–18.

33. Kim JH, Lane WS, Reinberg D. Human Elongator facilitates RNA polymerase II transcription through chromatin. Proc Natl Acad Sci U S A. 2002;99(3):1241–6.

34. Wittschieben BO, Otero G, de Bizemont T, Fellows J, Erdjument-Bromage H, Ohba R, et al. A novel histone acetyltransferase is an integral subunit of elongating RNA polymerase II holoenzyme. Mol Cell. 1999;4(1):123–8.

35. Dauden MI, Jaciuk M, Weis F, Lin TY, Kleindienst C, Abbassi NEH, et al. Molecular basis of tRNA recognition by the Elongator complex. Sci Adv. 2019;5(7):eaaw2326.

36. Glatt S, Séraphin B, Müller CW. Elongator: transcriptional or translational regulator? Transcription. 2012;3(6):273–6.

37. Lin TY, Abbassi NEH, Zakrzewski K, Chramiec-Głąbik A, Jemioła-Rzemińska M, Różycki J, et al. The Elongator subunit Elp3 is a non-canonical tRNA acetyltransferase. Nat Commun. 2019;10(1):625.

38. Goffena J, Lefcort F, Zhang Y, Lehrmann E, Chaverra M, Felig J, et al. Elongator and codon bias regulate protein levels in mammalian peripheral neurons. Nat Commun. 2018;9(1):889.

39. Norcliffe-Kaufmann L, Kaufmann H. Familial dysautonomia (Riley-Day syndrome): when baroreceptor feedback fails. Auton Neurosci. 2012;172(1-2):26–30.

40. Norcliffe-Kaufmann L, Slaugenhaupt SA, Kaufmann H. Familial dysautonomia: History, genotype, phenotype and translational research. Prog Neurobiol. 2017;152:131–48.

41. Pearson J, Pytel BA, Grover-Johnson N, Axelrod F, Dancis J. Quantitative studies of dorsal root ganglia and neuropathologic observations on spinal cords in familial dysautonomia. J Neurol Sci. 1978;35(1):77–92.

42. Gutierrez JV, Norcliffe-Kaufmann L, Kaufmann H. Brainstem reflexes in patients with familial dysautonomia. Clin Neurophysiol. 2015;126(3):626–33.

43. Norcliffe-Kaufmann L, Kaufmann H. Familial dysautonomia (Riley–Day syndrome). Primer on the Autonomic Nervous System: Elsevier; 2023. p. 527–31.

44. Morini E, Gao D, Montgomery CM, Salani M, Mazzasette C, Krussig TA, et al. ELP1 Splicing Correction Reverses Proprioceptive Sensory Loss in Familial Dysautonomia. Am J Hum Genet. 2019;104(4):638–50.

45. Cascella M, Muzio MR, Monaco F, Nocerino D, Ottaiano A, Perri F, et al. Pathophysiology of nociception and rare genetic disorders with increased pain threshold or pain insensitivity. Pathophysiology. 2022;29(3):435–52.

46. Axelrod FB, Iyer K, Fish I, Pearson J, Sein ME, Spielholz N. Progressive sensory loss in familial dysautonomia. Pediatrics. 1981;67(4):517–22.

47. González-Duarte A, Cotrina-Vidal M, Kaufmann H, Norcliffe-Kaufmann L. Familial dysautonomia. Clinical Autonomic Research. 2023:1–12.

48. Morini E, Chekuri A, Logan EM, Bolduc JM, Kirchner EG, Salani M, et al. Development of an oral treatment that rescues gait ataxia and retinal degeneration in a phenotypic mouse model of familial dysautonomia. The American Journal of Human Genetics. 2023;110(3):531–47.

49. Abashidze A, Gold V, Anavi Y, Greenspan H, Weil M. Involvement of IKAP in peripheral target innervation and in specific JNK and NGF signaling in developing PNS neurons. PLoS One. 2014;9(11):e113428.

50. George L, Chaverra M, Wolfe L, Thorne J, Close-Davis M, Eibs A, et al. Familial dysautonomia model reveals Ikbkap deletion causes apoptosis of Pax3+ progenitors and peripheral neurons. Proc Natl Acad Sci U S A. 2013;110(46):18698–703.

51. Lee G, Papapetrou EP, Kim H, Chambers SM, Tomishima MJ, Fasano CA, et al. Modelling pathogenesis and treatment of familial dysautonomia using patient-specific iPSCs. Nature. 2009;461(7262):402-6.

52. Morini E, Dietrich P, Salani M, Downs HM, Wojtkiewicz GR, Alli S, et al. Sensory and autonomic deficits in a new humanized mouse model of familial dysautonomia. Hum Mol Genet. 2016;25(6):1116–28.

53. Chekuri A, Logan EM, Krauson AJ, Salani M, Ackerman S, Kirchner EG, et al. Selective retinal ganglion cell loss and optic neuropathy in a humanized mouse model of familial dysautonomia. Hum Mol Genet. 2021.

54. Okaty BW, Commons KG, Dymecki SM. Embracing diversity in the 5-HT neuronal system. Nat Rev Neurosci. 2019;20(7):397–424.

55. Fozard JR. Neuronal 5-HT receptors in the periphery. Neuropharmacology. 1984;23(12b):1473–86.

56. Maurer P, T’Sas F, Coutte L, Callens N, Brenner C, Van Lint C, et al. FEV acts as a transcriptional repressor through its DNA-binding ETS domain and alanine-rich domain. Oncogene. 2003;22(21):3319–29.

57. McLean PG, Borman RA, Lee K. 5-HT in the enteric nervous system: gut function and neuropharmacology. Trends Neurosci. 2007;30(1):9–13.

58. Watanabe H, Shimohigashi M, Yokohari F. Serotonin-immunoreactive sensory neurons in the antenna of the cockroach Periplaneta americana. J Comp Neurol. 2014;522(2):414–34.

59. Wyler SC, Spencer WC, Green NH, Rood BD, Crawford L, Craige C, et al. Pet-1 Switches Transcriptional Targets Postnatally to Regulate Maturation of Serotonin Neuron Excitability. J Neurosci. 2016;36(5):1758–74.

60. Nichols RC, Blinder J, Pai SI, Ge Q, Targoff IN, Plotz PH, et al. Assignment of two human autoantigen genes-isoleucyl-tRNA synthetase locates to 9q21 and lysyl-tRNA synthetase locates to 16q23-q24. Genomics. 1996;36(1):210–3.

61. Heng HH, Shi XM, Scherer SW, Andrulis IL, Tsui LC. Refined localization of the asparagine synthetase gene (ASNS) to chromosome 7, region q21.3, and characterization of the somatic cell hybrid line 4AF/106/KO15. Cytogenet Cell Genet. 1994;66(2):135–8.

62. Liu G, Maunoury C, Kamoun P, Aral B. Assignment of the human gene encoding the delta 1-pyrroline-5-carboxylate synthetase (P5CS) to 10q24.3 by in situ hybridization. Genomics. 1996;37(1):145–6.

63. Vékony N, Wolf S, Boissel JP, Gnauert K, Closs EI. Human cationic amino acid transporter hCAT-3 is preferentially expressed in peripheral tissues. Biochemistry. 2001;40(41):12387–94.

64. Rebsamen M, Girardi E, Sedlyarov V, Scorzoni S, Papakostas K, Vollert M, et al. Gain-of-function genetic screens in human cells identify SLC transporters overcoming environmental nutrient restrictions. Life Sci Alliance. 2022;5(11).

65. Cruzen ME, Bengtsson U, McMahon J, Wasmuth JJ, Arfin SM. Assignment of the cysteinyl-tRNA synthetase gene (CARS) to 11p15.5. Genomics. 1993;15(3):692–3.

66. Tavares-Ferreira D, Shiers S, Ray PR, Wangzhou A, Jeevakumar V, Sankaranarayanan I, et al. Spatial transcriptomics of dorsal root ganglia identifies molecular signatures of human nociceptors. Sci Transl Med. 2022;14(632):eabj8186.

67. Usoskin D, Furlan A, Islam S, Abdo H, Lönnerberg P, Lou D, et al. Unbiased classification of sensory neuron types by large-scale single-cell RNA sequencing. Nat Neurosci. 2015;18(1):145–53.

68. Yang L, Xu M, Bhuiyan SA, Li J, Zhao J, Cohrs RJ, et al. Human and mouse trigeminal ganglia cell atlas implicates multiple cell types in migraine. Neuron. 2022;110(11):1806–21.e8.

69. Tavares-Ferreira D, Shiers S, Ray PR, Wangzhou A, Jeevakumar V, Sankaranarayanan I, et al. Spatial transcriptomics of dorsal root ganglia identifies molecular signatures of human nociceptors. Science translational medicine. 2022;14(632):eabj8186.

70. Wu HF, Yu W, Saito-Diaz K, Huang CW, Carey J, Lefcort F, et al. Norepinephrine transporter defects lead to sympathetic hyperactivity in Familial Dysautonomia models. Nat Commun. 2022;13(1):7032.

71. Tolman Z, Chaverra M, George L, Lefcort F. Elp1 is required for development of visceral sensory peripheral and central circuitry. bioRxiv. 2022:2022.04. 11.487663.

72. Leonard CE, Quiros J, Lefcort F, Taneyhill LA. Loss of Elp1 disrupts trigeminal ganglion neurodevelopment in a model of familial dysautonomia. Elife. 2022;11:e71455.

73. Narasimhan VM, Hunt KA, Mason D, Baker CL, Karczewski KJ, Barnes MR, et al. Health and population effects of rare gene knockouts in adult humans with related parents. Science. 2016;352(6284):474–7.

74. Kojic M, Abbassi NEH, Lin TY, Jones A, Wakeling EL, Clement E, et al. A novel ELP1 mutation impairs the function of the Elongator complex and causes a severe neurodevelopmental phenotype. J Hum Genet. 2023.

75. Hims MM, Shetty RS, Pickel J, Mull J, Leyne M, Liu L, et al. A humanized IKBKAP transgenic mouse models a tissue-specific human splicing defect. Genomics. 2007;90(3):389–96.

76. Dietrich P, Alli S, Shanmugasundaram R, Dragatsis I. IKAP expression levels modulate disease severity in a mouse model of familial dysautonomia. Hum Mol Genet. 2012;21(23):5078–90.

77. Dietrich P, Yue J, E S, Dragatsis I. Deletion of exon 20 of the Familial Dysautonomia gene Ikbkap in mice causes developmental delay, cardiovascular defects, and early embryonic lethality. PLoS One. 2011;6(10):e27015.

78. Dobin A, Davis CA, Schlesinger F, Drenkow J, Zaleski C, Jha S, et al. STAR: ultrafast universal RNA-seq aligner. Bioinformatics. 2013;29(1):15–21.

79. Bolger AM, Lohse M, Usadel B. Trimmomatic: a flexible trimmer for Illumina sequence data. Bioinformatics. 2014;30(15):2114–20.

80. Anders S, Pyl PT, Huber W. HTSeq--a Python framework to work with high-throughput sequencing data. Bioinformatics. 2015;31(2):166–9.

81. Love MI, Huber W, Anders S. Moderated estimation of fold change and dispersion for RNA-seq data with DESeq2. Genome Biol. 2014;15(12):550.

82. Leek JT. svaseq: removing batch effects and other unwanted noise from sequencing data. Nucleic Acids Res. 2014;42(21):e161.

83. Donadon I, Pinotti M, Rajkowska K, Pianigiani G, Barbon E, Morini E, et al. Exon-specific U1 snRNAs improve ELP1 exon 20 definition and rescue ELP1 protein expression in a familial dysautonomia mouse model. Human molecular genetics. 2018;27(14):2466–76.

84. Doncheva NT, Morris JH, Gorodkin J, Jensen LJ. Cytoscape StringApp: Network Analysis and Visualization of Proteomics Data. J Proteome Res. 2019;18(2):623–32.

85. Shannon P, Markiel A, Ozier O, Baliga NS, Wang JT, Ramage D, et al. Cytoscape: a software environment for integrated models of biomolecular interaction networks. Genome Res. 2003;13(11):2498–504.

86. Subramanian A, Tamayo P, Mootha VK, Mukherjee S, Ebert BL, Gillette MA, et al. Gene set enrichment analysis: a knowledge-based approach for interpreting genome-wide expression profiles. Proc Natl Acad Sci U S A. 2005;102(43):15545–50.

87. Langfelder P, Horvath S. WGCNA: an R package for weighted correlation network analysis. BMC Bioinformatics. 2008;9:559.

88. Cortal A, Martignetti L, Six E, Rausell A. Gene signature extraction and cell identity recognition at the single-cell level with Cell-ID. Nat Biotechnol. 2021;39(9):1095–102.

89. Hao Y, Hao S, Andersen-Nissen E, Mauck WM, 3rd, Zheng S, Butler A, et al. Integrated analysis of multimodal single-cell data. Cell. 2021;184(13):3573–87.e29.

90. Choudhary S, Satija R. Comparison and evaluation of statistical error models for scRNA-seq. Genome Biol. 2022;23(1):27.

