## Supplementary and Supporting Figures with Legends for "Transcriptome analysis in a humanized mouse model of familial dysautonomia reveals tissue-specific gene expression disruption in the peripheral nervous system"

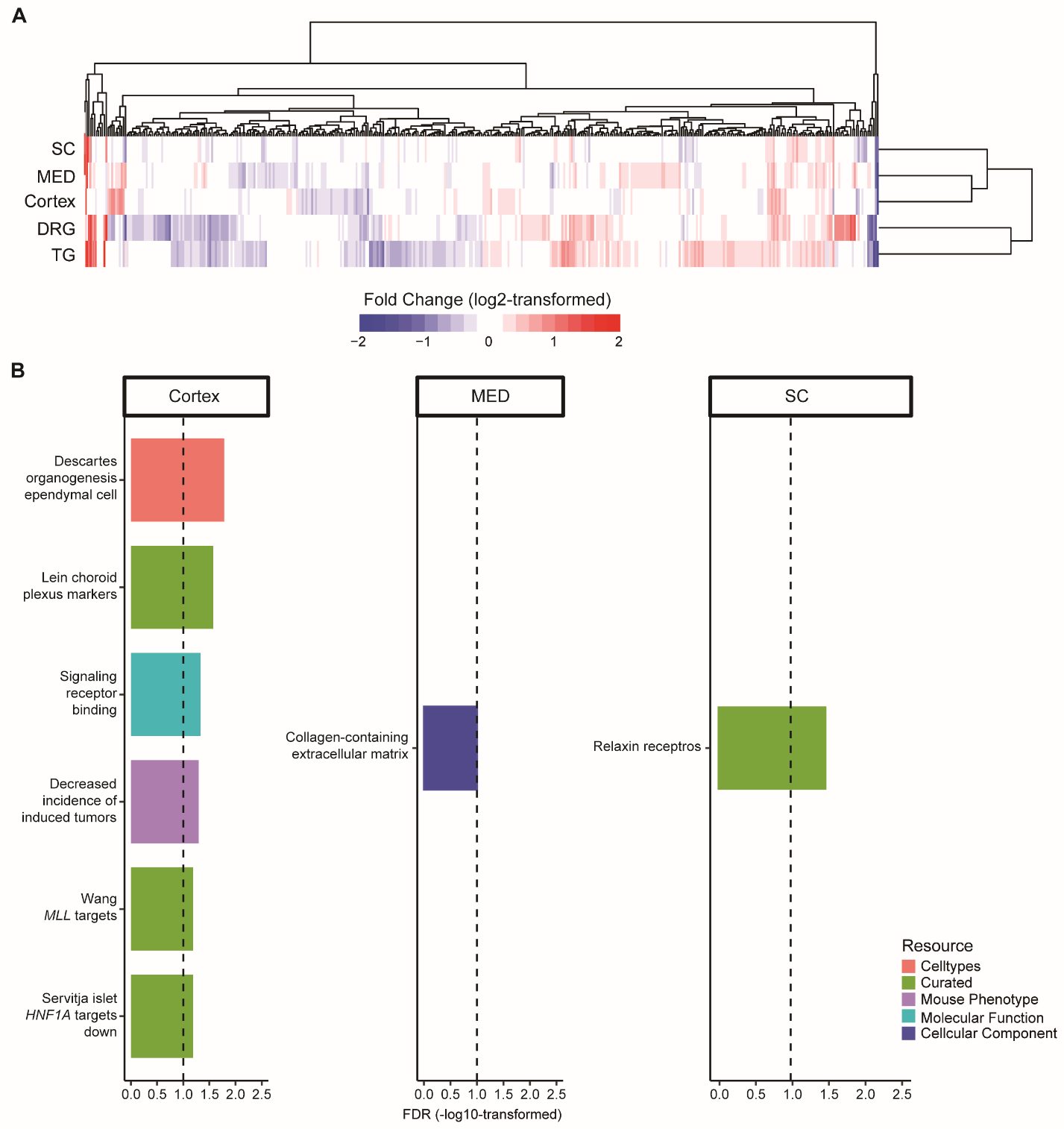

**Supplementary Figure S1. Tissue-specific DEGs and their functional enrichment.**

**(A)** The heatmap represents all the DEGs across five tissues. Each column represents a gene, and each row represents a tissue. The red color domain represents upregulation between FD and Control while the blue color domain represents downregulation, where the expression changes are measured by log2-transformed fold changes. The deeper the color, the stronger the fold change. **(B)** The bar plot represents the functional enrichment based on DEGs in Cortex, MED, and SC, respectively. Note, all significant terms (FDR < 0.1) are displayed per tissue. The x-axis represents the enrichment significance in -log10-transformed FDR while the y-axis represents the terms. The bar colors indicate the resources of Gene Set Enrichment Analysis (GSEA) v. MS1. The vertical black dashed line represents an FDR of 0.1.

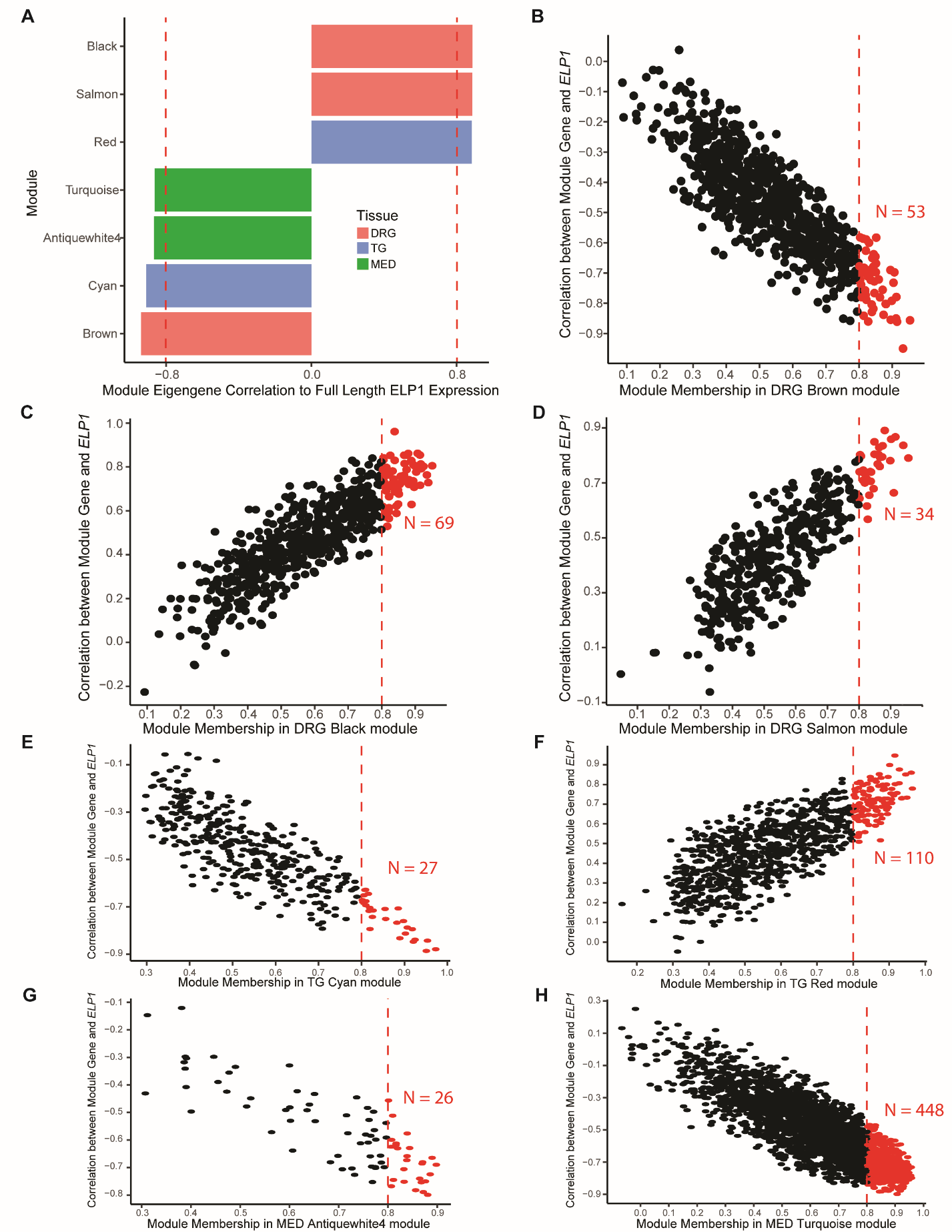

**Supplementary Figure S2. Identification of *ELP1* dose-responsive genes.**

**(A)** The bar plot represents how strong the full-length *ELP1* expression is correlated with the eigengene of each tissue-specific co-expression module. Only modules with an absolute Pearson correlation coefficient no less than 0.8 are displayed. The x-axis demonstrates the Pearson correlation coefficient between the full-length *ELP1* expression and an eigengene. The y-axis indicates the module names. The bars are colored by tissue types. The vertical dashed red line indicates Pearson correlation coefficients at -0.8 and 0.8, respectively. **(B-H)** The scatter plots represent the relationships among each gene in the modules in panel **(A)**, module eigengenes, and the full-length *ELP1* expression. The x-axis demonstrates the module membership, which is the Pearson correlation coefficient between each gene in the module and the module’s eigengene. The y-axis demonstrates the Pearson correlation coefficient between each gene in the module and the full-length *ELP1* expression. The vertical dashed red line indicates a Pearson correlation coefficient cutoff at 0.8. The genes to the right side of this cutoff line are defined as *ELP1* dose-responsive genes and displayed in red dots. **(B)** The co-expression module of DRG Brown. **(C)** The co-expression module of DRG Black. **(D)** The co-expression module of DRG Salmon. **(E)** The co-expression module of TG Cyan. **(F)** The co-expression module of TG Red. **(G)** The co-expression module of MED Antiquewhite4. **(H)** The co-expression module of MED Turquoise.

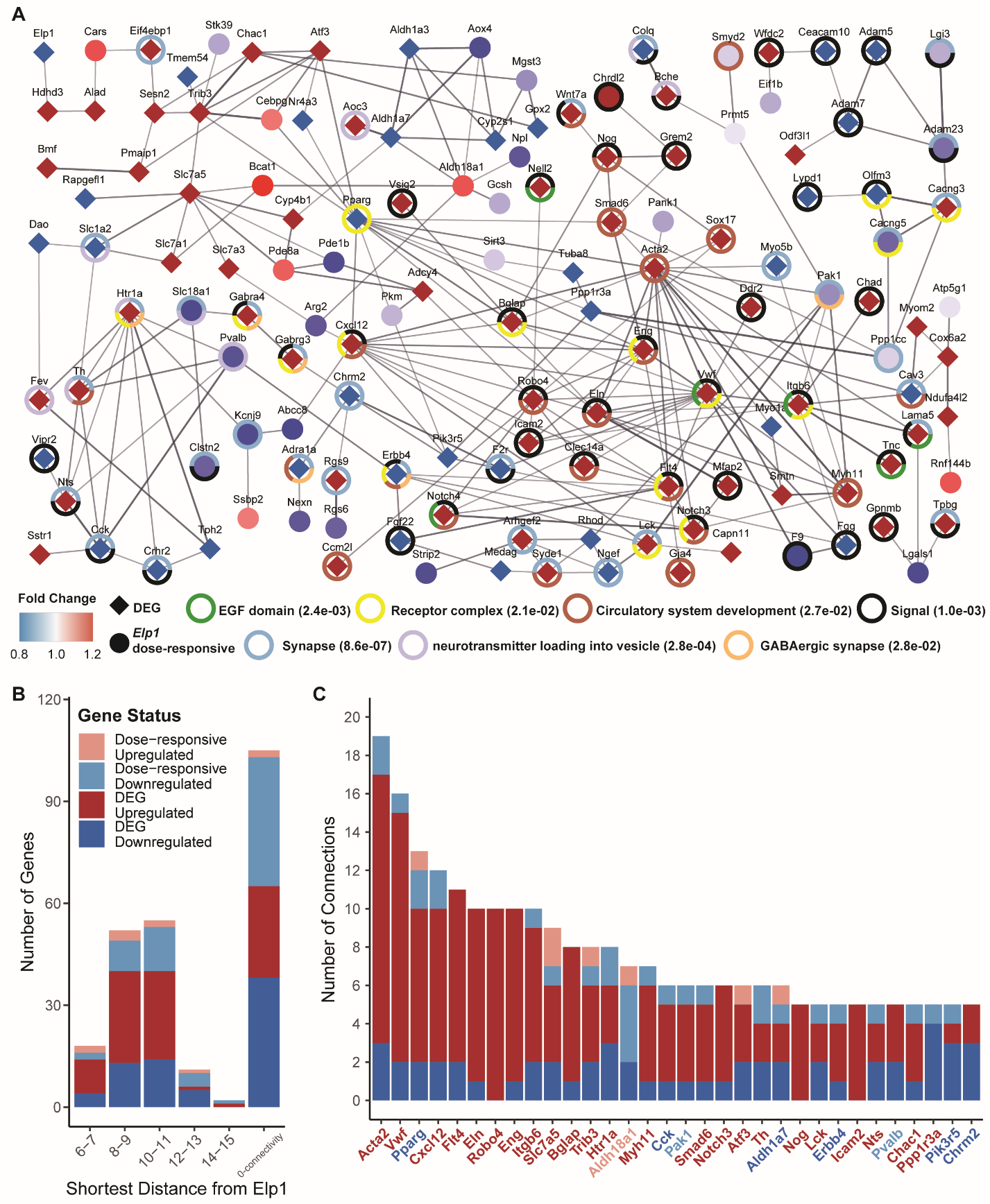

**Supplementary Figure S3. TG-specific dysregulated gene network due to ELP1 reduction.**

**(A)** The dysregulated gene network in TG. Each node is either a DEG indicated by a diamond shape or an ELP1 dose-responsive gene indicated by a round shape. The colors for the nodes reflect the fold changes in the genes between FD and Control. The red color domain represents upregulation between FD and Control while the blue color domain represents downregulation. The deeper the color, the stronger the fold changes. Each edge represents a potential interaction between the two connected genes, where only an interaction score of more than 0.4 (default) in String-DB is displayed. The thicker the edge, the higher the interaction score. Only the dysregulated genes with at least one interaction are displayed. The rings outside the nodes represent significant functional enrichment with FDR < 0.1 using all the dysregulated genes (i.e., DEGs and ELP1 dose-responsive genes). The associated functional enrichment terms with the ring colors are given, where the values in the brackets are the enrichment FDR values for the terms. **(B)** The bar plot demonstrates the number of dysregulated genes in TG at different distances to ELP1. The x-axis represents the distance of the shortest path to a gene. The genes in the “0-connectivity” distance category refer to those dysregulated genes not displayed in panel **(A)** due to an interaction score >= 0.4. The y-axis represents the number of genes at each distance. **(C)** The bar plot demonstrates hub genes in TG ranked by their number of connections to the neighbor genes in the network of panel **(A)**. The x-axis represents the hub gene names, where each gene is colored according to its dysregulation direction and gene category.

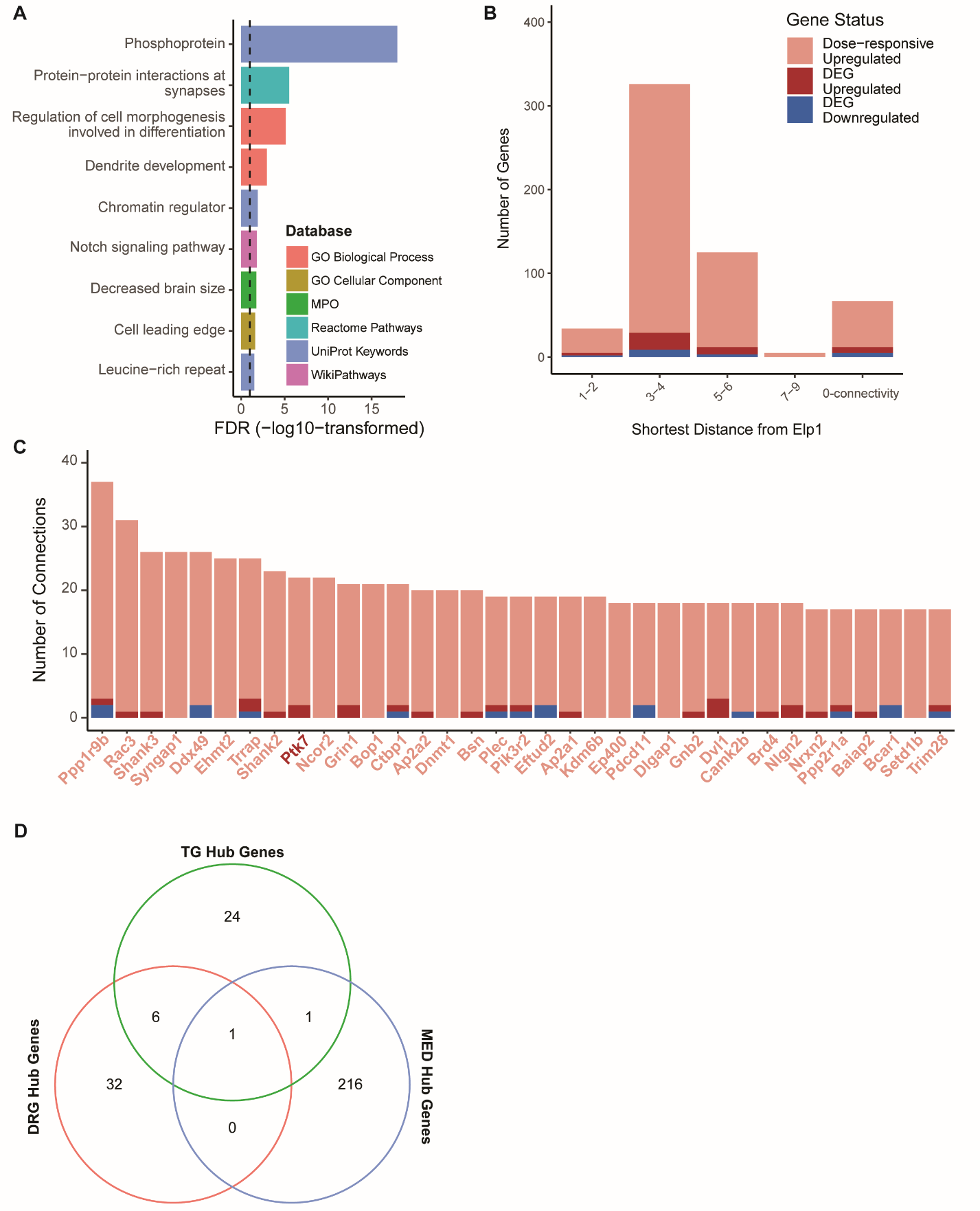

**Supplementary Figure S4. MED-specific dysregulated gene network due to ELP1 reduction.**

**(A)** The bar plot represents the GO functional enrichment term with FDR < 0.1 using all the dysregulated genes (i.e., DEGs and ELP1 dose-responsive genes). The x-axis demonstrates the FDR value of enrichment in the -log10-transformed scale. The y-axis demonstrates the GO terms where bars are colored by GO resources. The vertical dashed black line indicates an FDR of 0.1. **(B)** The bar plot demonstrates the number of dysregulated genes in MED at different distances to ELP1. The x-axis represents the distance of the shortest path to a gene. The genes in the “0-connectivity” distance category refer to those dysregulated genes without any connected path to ELP1. The y-axis represents the number of genes at each distance. **(C)** The bar plot demonstrates hub genes in MED ranked by their number of connections to the neighbor genes in the network of panel **(A)**. The x-axis represents the hub gene names, where each gene is colored according to its dysregulation direction and gene category. **(D)** The Venn diagram shows the overlaps among the hub genes of the dysregulated gene networks in DRG, TG, and MED. The numbers represent the amount of overlapped genes.

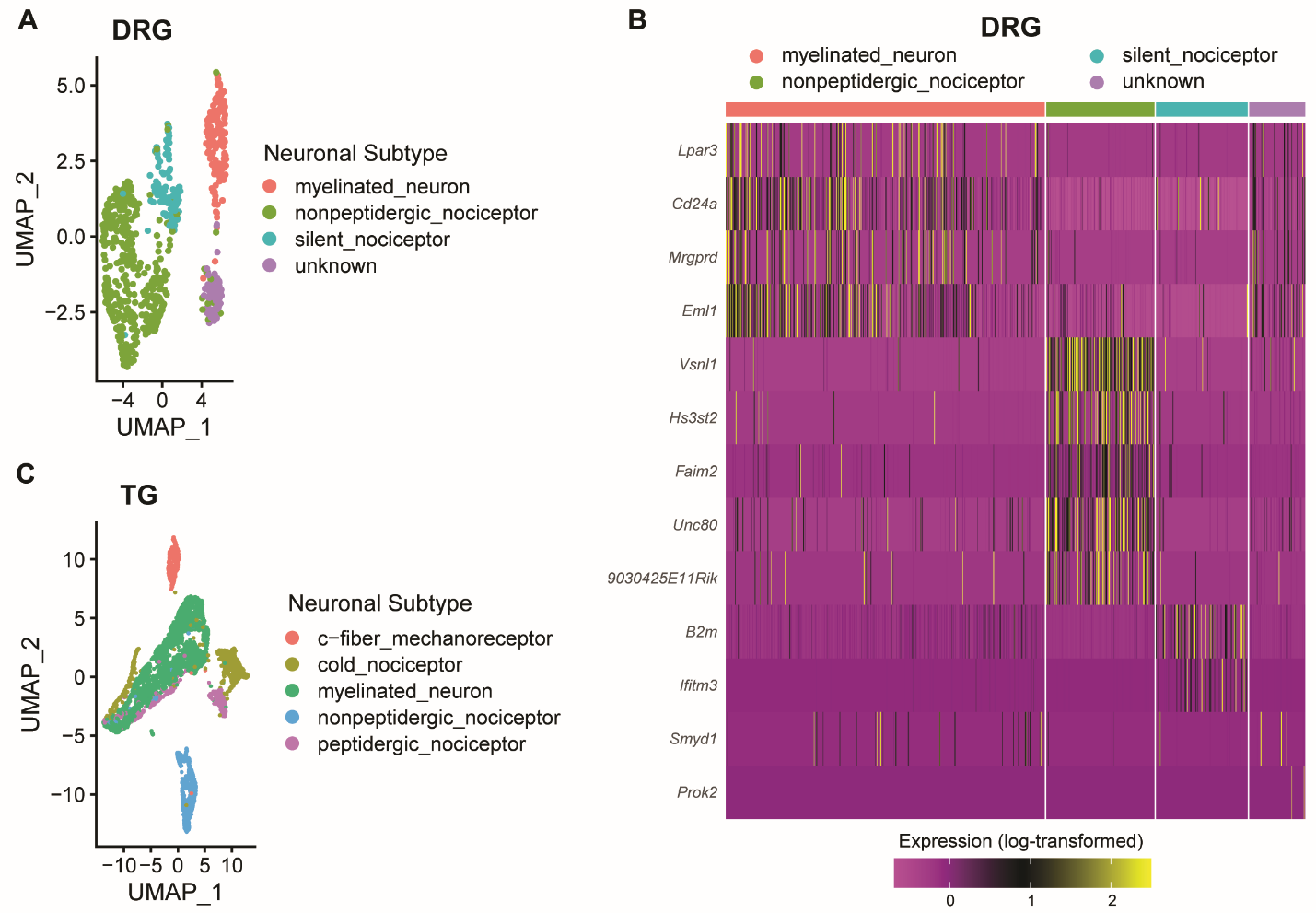

**Supplementary Figure S5. Re-analyses of publicly available scRNA datasets of mouse DRG and TG.**

**(A)** The scatter plot represents the 2D UMAP space of mouse DRG. Each dot represents a cell and is colored according to its assigned neuronal subtype from the re-analysis. **(B)** The heatmap represents the expression of selected markers of each neuronal subtype in mouse DRG. The rows represent neuronal subtype markers while the columns represent cells grouped by their assigned neuronal subtypes from the re-analysis. The gene expression is measured in the log-transformed scale and colored. The purple domain reflects lower expression while the yellow domain represents higher expression. **(C)** The scatter plot represents the 2D UMAP space of mouse TG. Each dot represents a cell and is colored according to its assigned neuronal subtype from the re-analysis.
